## supplementary material for "Does ankle push-off correct for errors in anterior-posterior foot placement relative to center-of-mass states?"

Content:

S1:	The regression coefficients of the CoM states in the linear foot placement model and their statistical significance in normal and slow walking;

S2:	SPM regression test of the foot placement error and the combined AP GRF for every participant in normal and slow walking;

S3:	SPM regression test of the foot placement error and the trailing leg’s AP GRF for every participant in normal and slow walking;

S4:	SPM regression test of the foot placement error and the ankle moment for every participant in normal and slow walking;

S5:	Mean kinetics, standard deviation, and group-level paired t-test of the kinetics time series (combined and trailing leg’s AP GRF, ankle moment) corresponding to the ‘largest’ and ‘smallest’ foot placement error in normal and slow walking.

## S1:

| (a) | (b) |
| --- | --- |
| 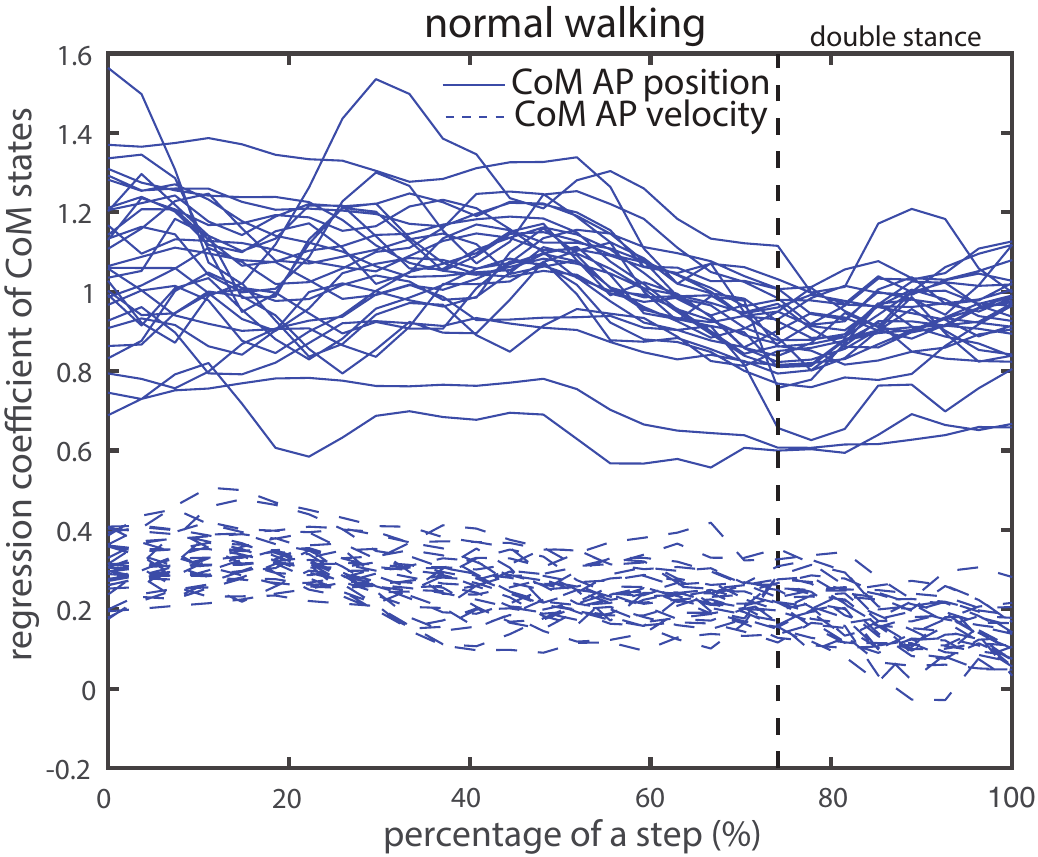 | 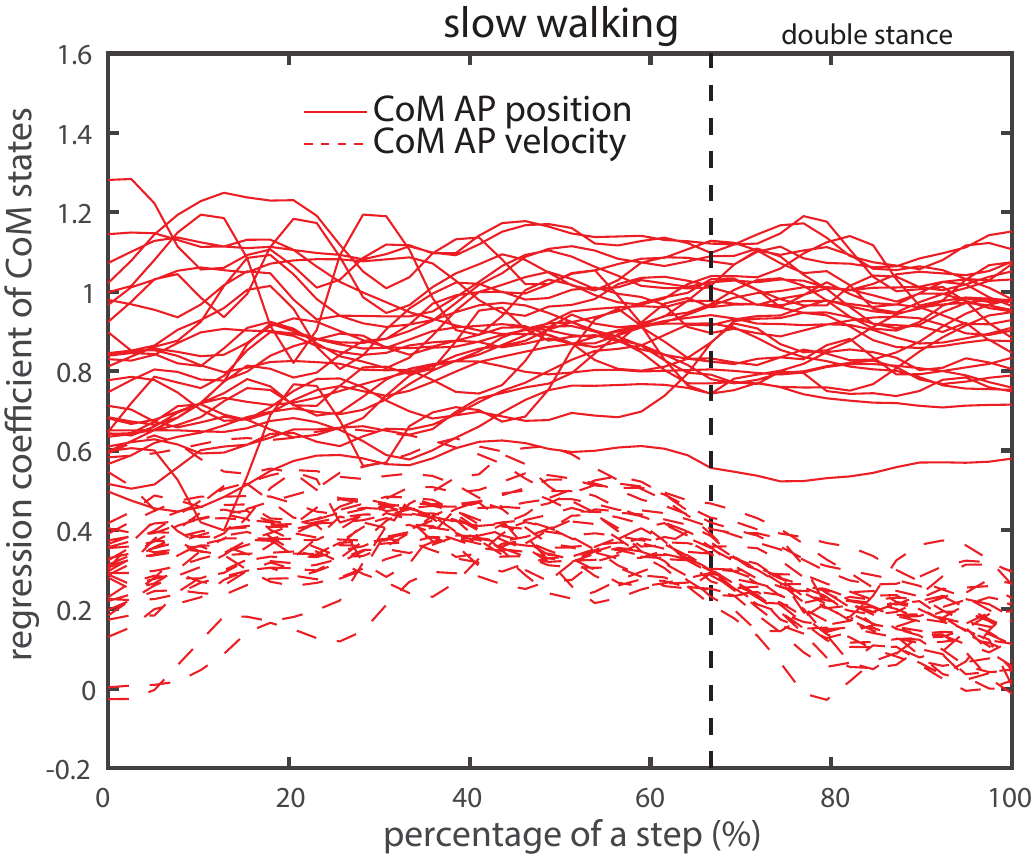 |
| (c) | (d) |
| 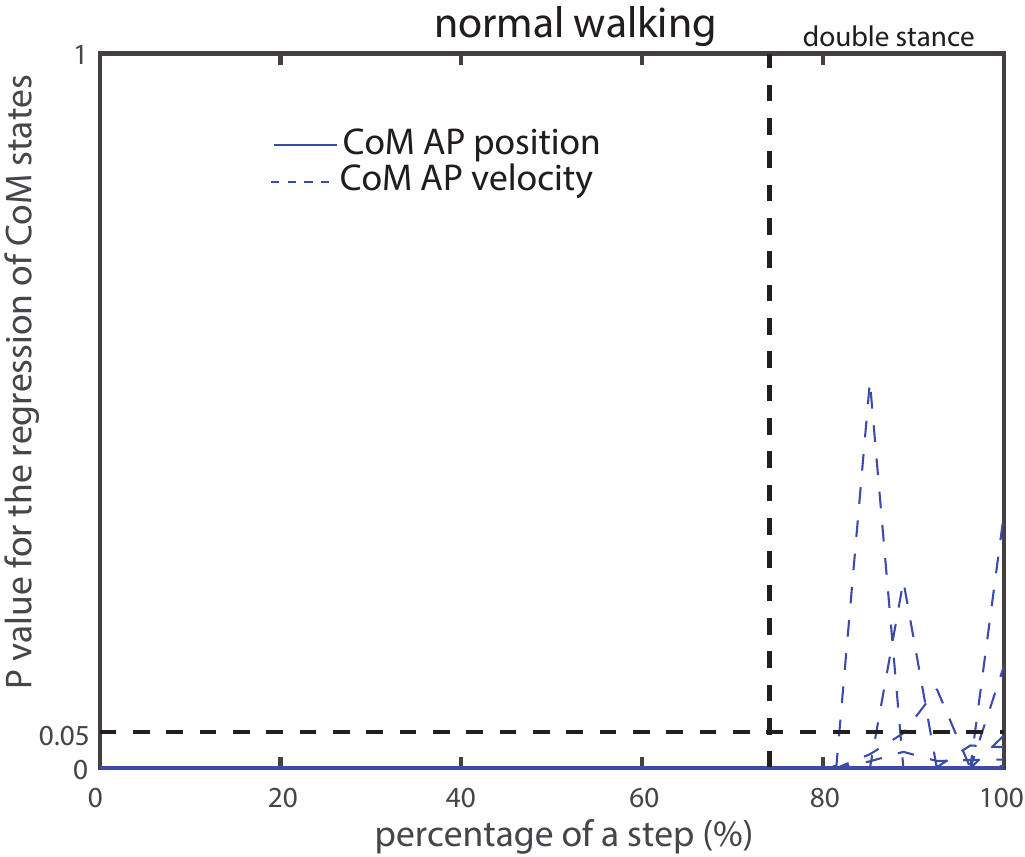 | 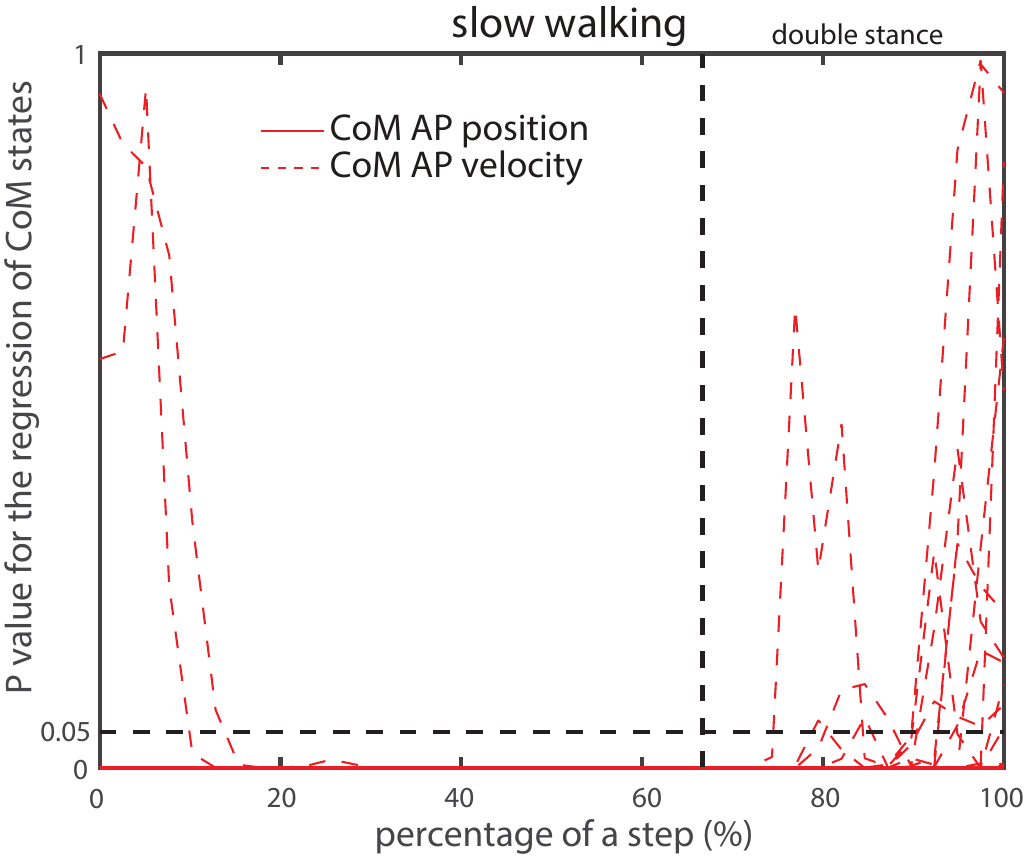 |

### Fig. S1 (a) Regression coefficients of the CoM states in the linear foot placement model in normal walking, and in (b) slow walking. The curves are the regression coefficients for each participant; (c/d) p-values of the linear regression of the CoM states in normal and slow walking. The CoM AP position is always significant as predictor of foot placement for all participants in normal and slow walking, while CoM AP velocity is significant for almost all participants in several phases at the beginning and end of a step in normal and slow walking.

## S2:

| (a) | 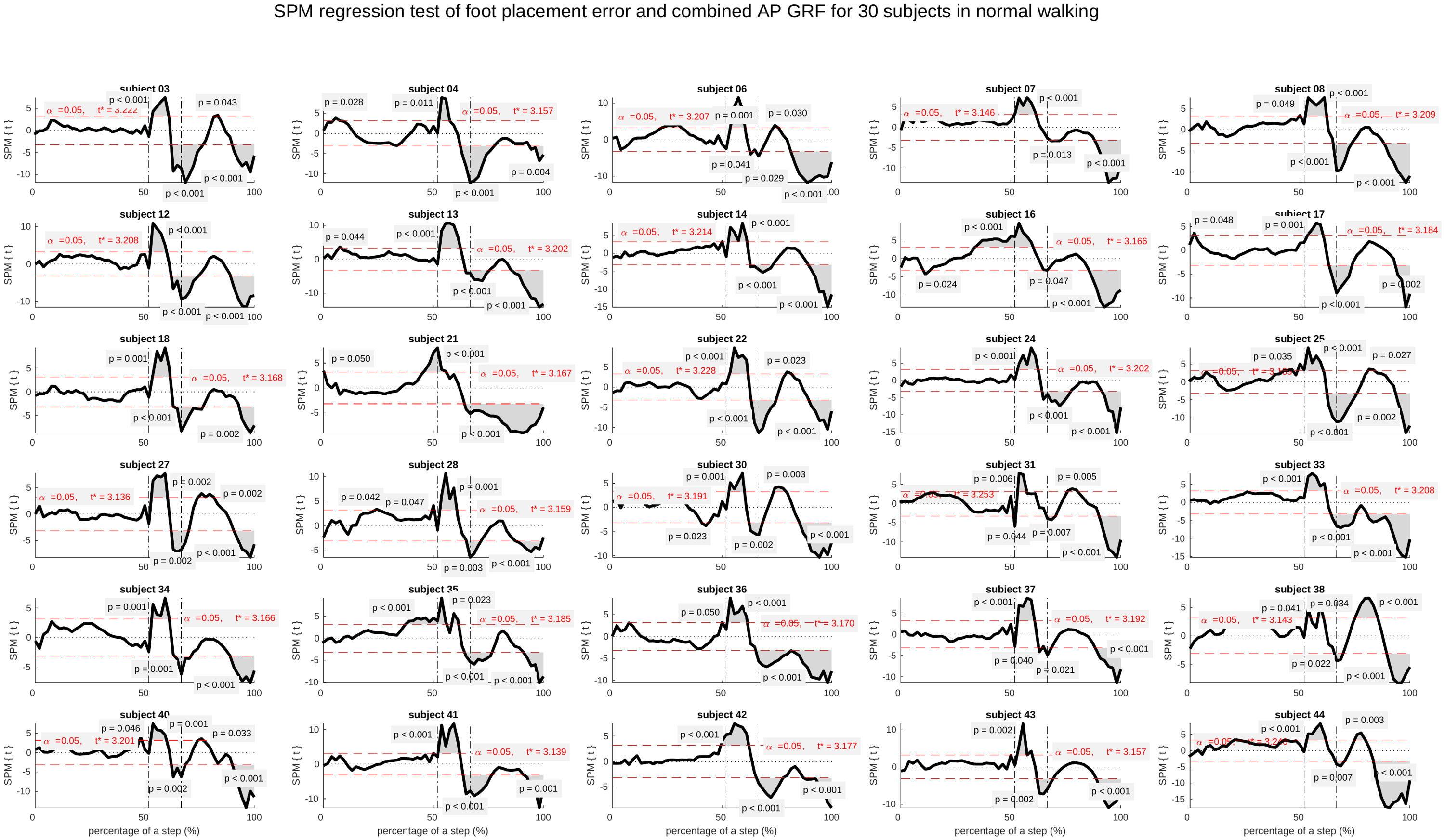 |
| --- | --- |
| (b) | 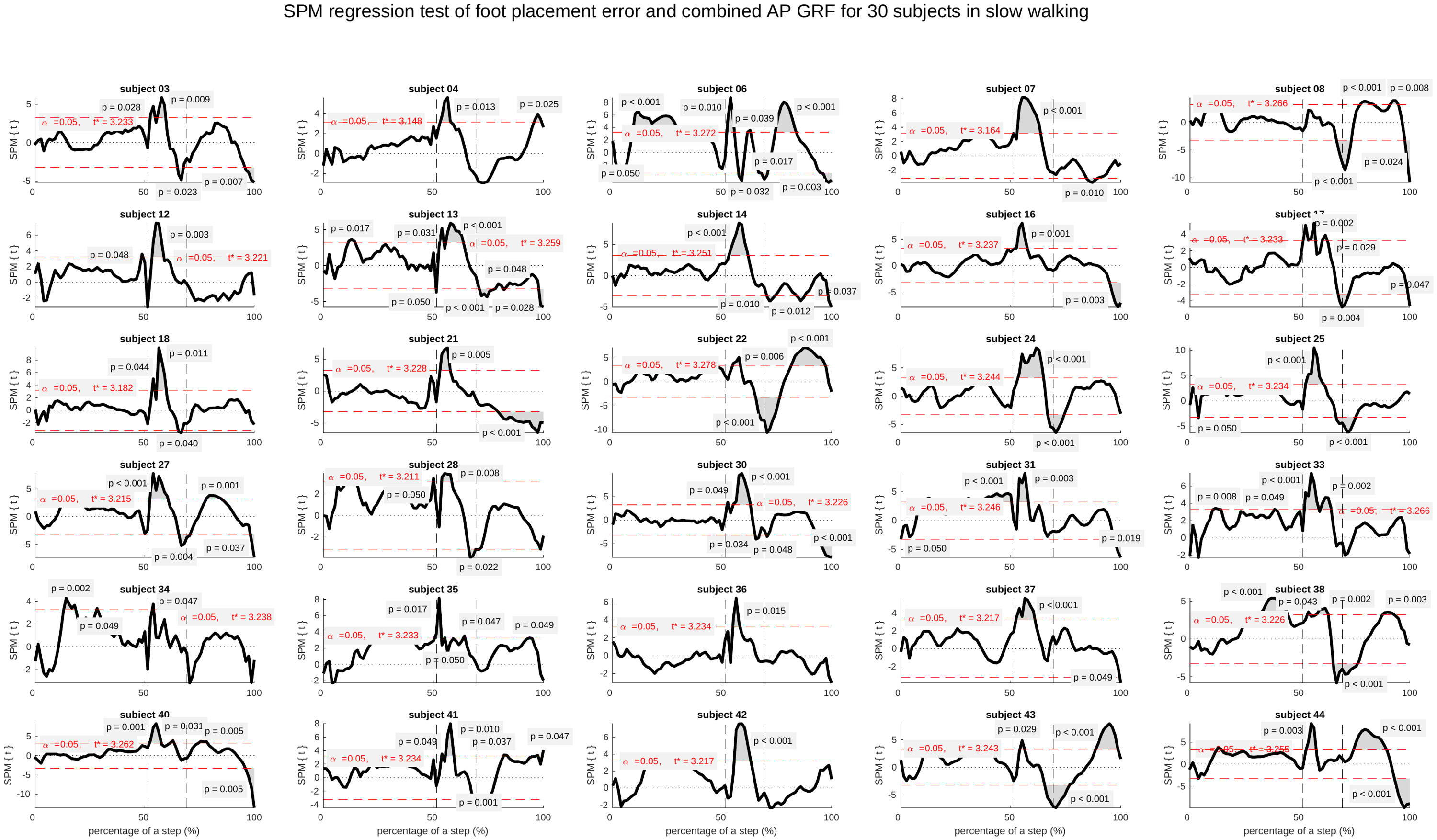 |

Fig. S2 (a) SPM regression test of the foot placement error and the combined AP GRF for 30 participants in normal walking. For all participants, early phases of the double stance are significant (p<0.05); (b) The same for slow walking. For all participants except for participant 8, early phases of the double stance are significant (p<0.05).

## S3:

| (a) | 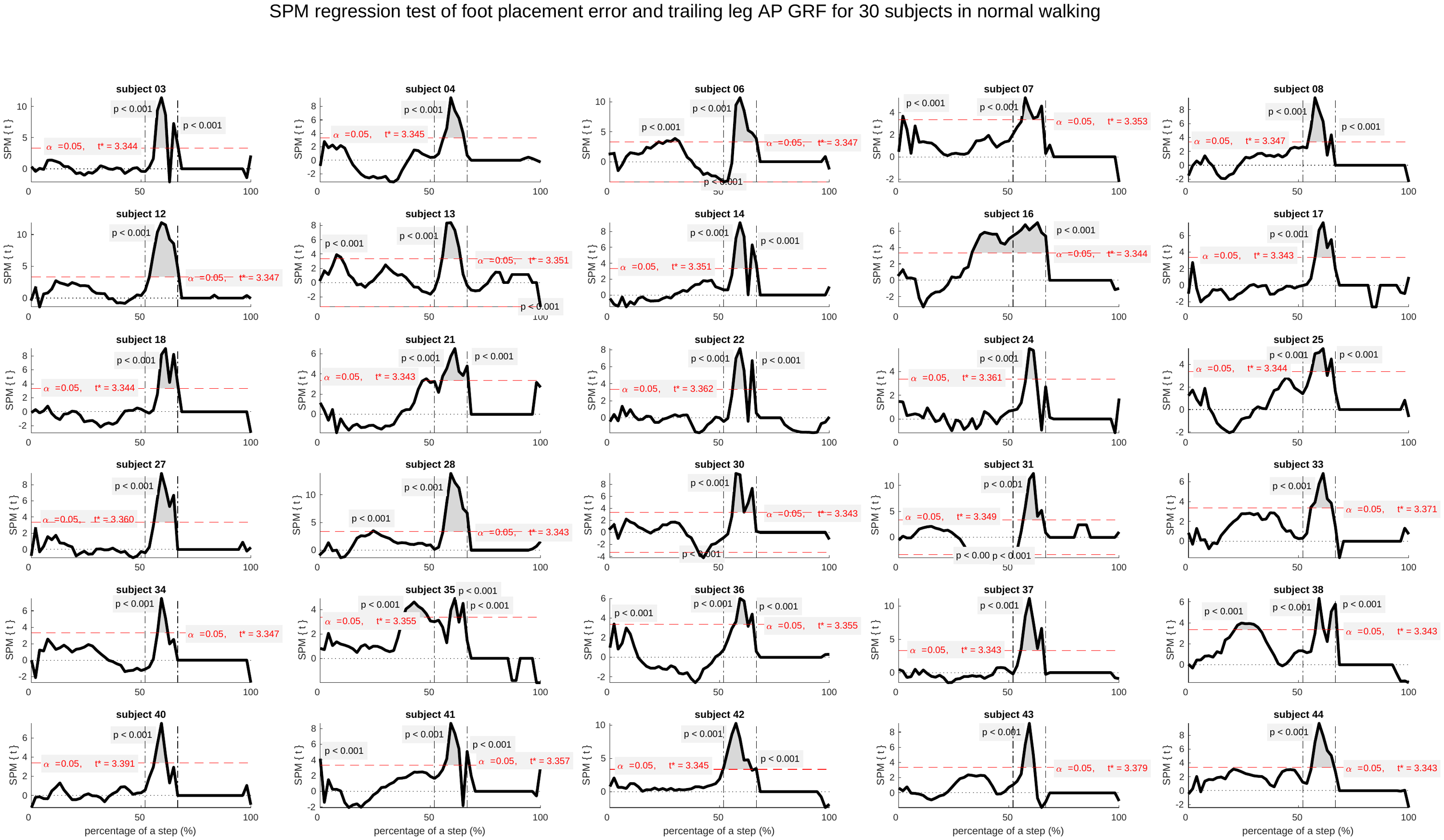 |
| --- | --- |
| (b) | 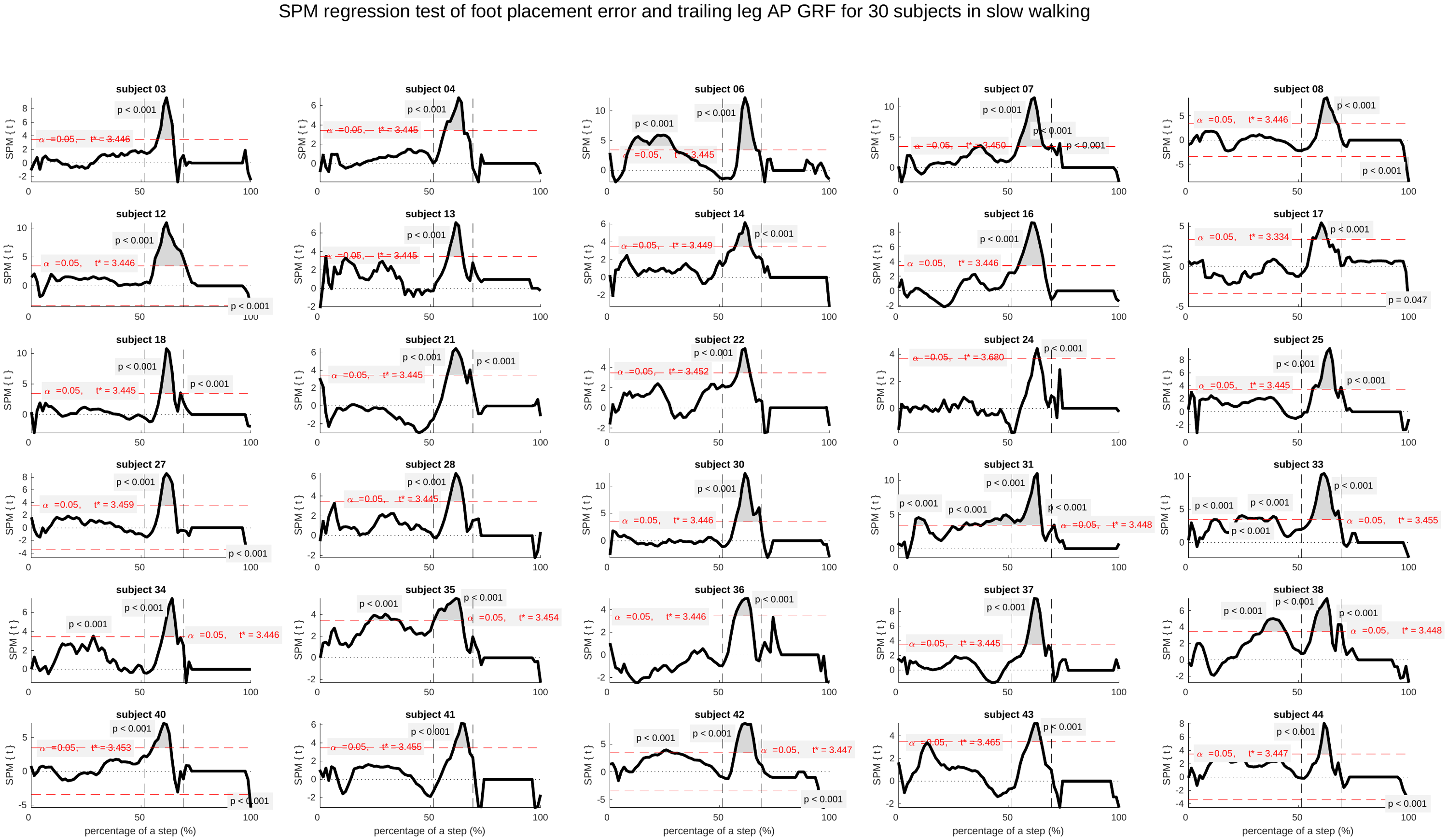 |

Fig. S3 SPM regression test of the foot placement error and the trailing leg’s AP GRF for 30 participants, (a) in normal walking, (b) in slow walking. For all participants, most of the double stances are significant (p<0.001).

## S4:

| (a) | 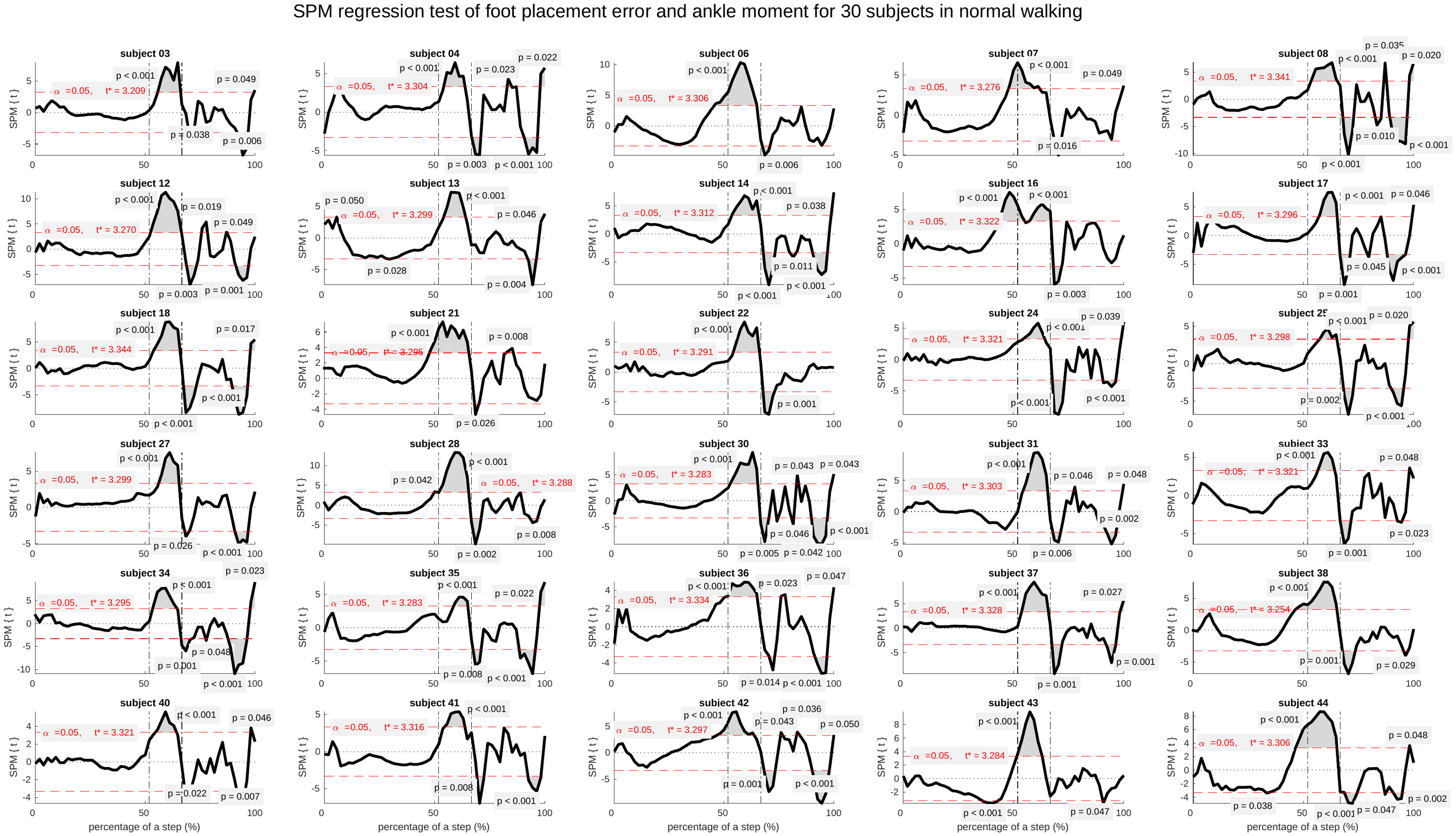 |
| --- | --- |
| (b) | 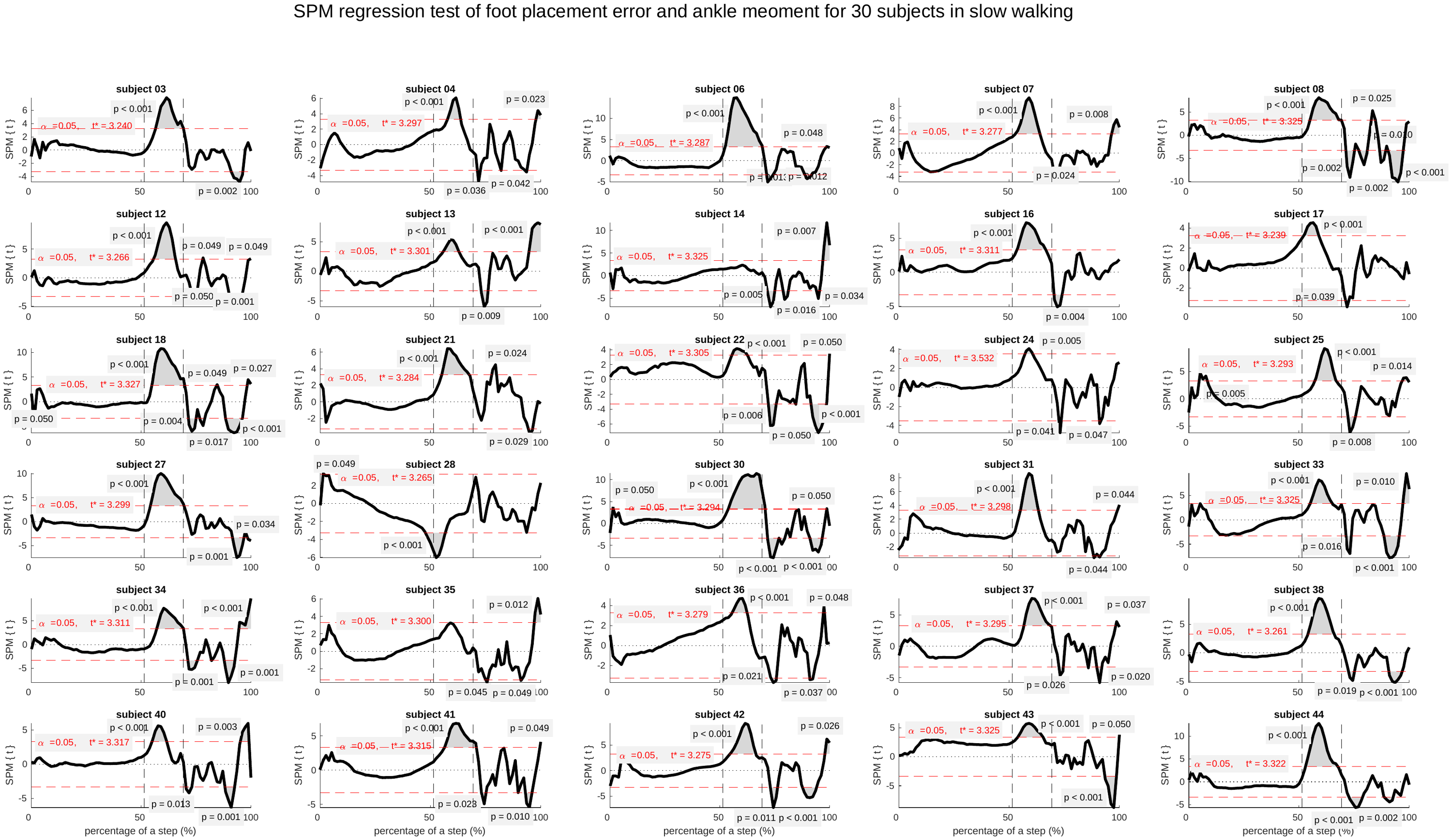 |

Fig. S4 (a) SPM regression test of the foot placement error and the ankle moment for 30 participants in normal walking. For all participants, the majority of the double stance is significant (p<0.001); (b) Also in slow walking, most of the double stances are significant (p<0.001) except for participant 14 and 35, the correlations during the double stance phase are not significant. For participant 28, the correlations during the double stance phase are negative, opposite to the expectations.

## S5:

Participant 24 had only 18 strides available in slow walking for analysis of the trailing leg’s AP GRF and the ankle moment because of stepping on both sides of the treadmill. The corresponding data were excluded for comparing trailing leg’s AP GRF and ankle moment for the “largest” and the “smallest” foot placement error.

| (a) | (b) |
| --- | --- |
| 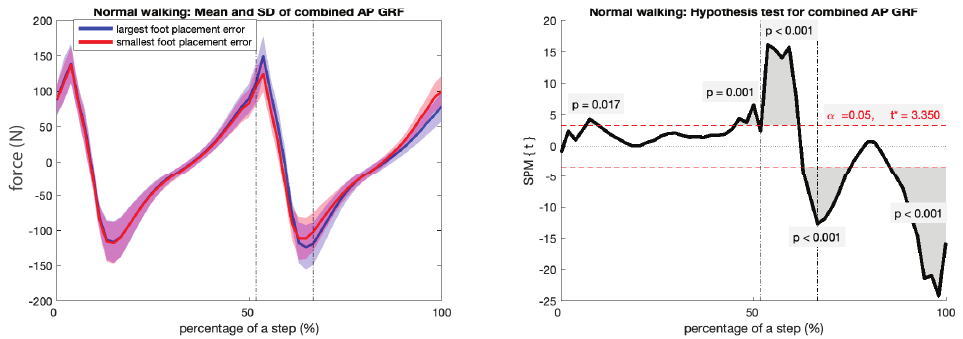 | |
| (c) | (d) |
| 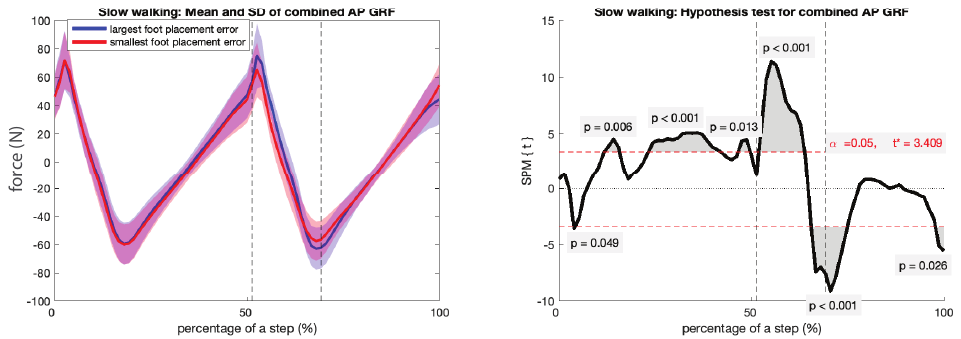 | |
| (e) | (f) |
| 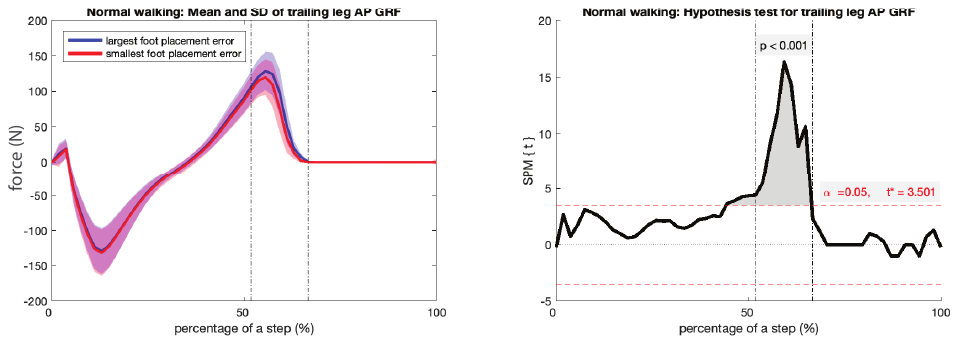 | |

| (g) | (h) |
| --- | --- |
| 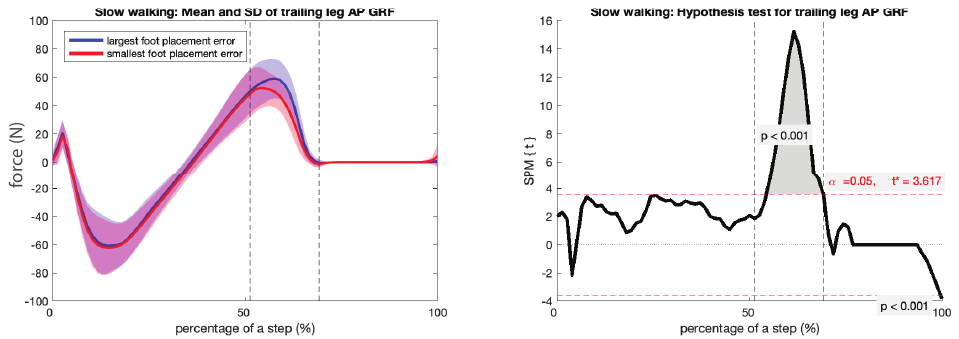 | |
| (i) | (j) |
| 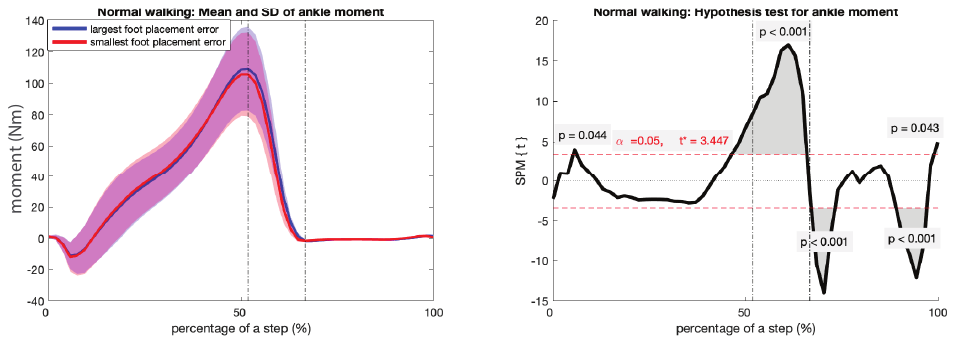 | |
| (k) | (l) |
| 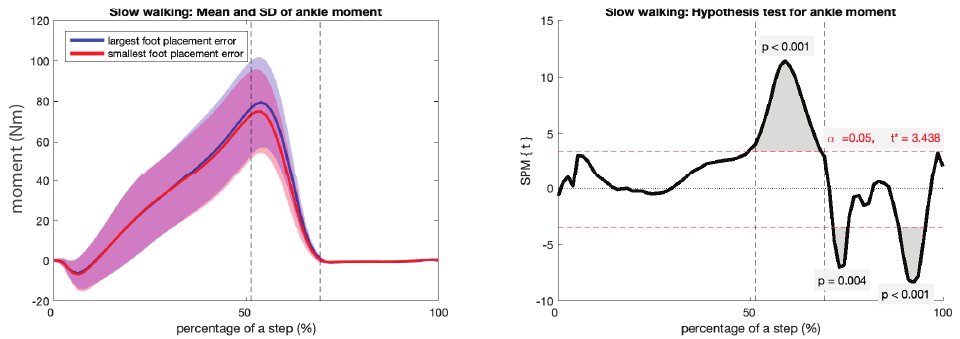 | |

Fig. S5 Mean kinetics, standard deviation (left figure) and group level paired t-test (right figure) for participant’s “largest” and “smallest” foot placement error: (a/b) combined AP GRF in normal walking; (c/d) combined AP GRF in slow walking; (e/f) trailing leg’s AP GRF in normal walking; (g/h) trailing leg’s AP GRF in slow walking; (i/j) ankle moment in normal walking; (k/l) ankle moment in slow walking.
